# Hydrophilic Shell Matrix Proteins of *Nautilus pompilius* and The Identification of a Core Set of Conchiferan Domains

**DOI:** 10.1101/2020.11.14.382804

**Authors:** Davin H. E. Setiamarga, Kazuki Hirota, Masa-aki Yoshida, Yusuke Takeda, Keiji Kito, Keisuke Shimizu, Yukinobu Isowa, Kazuho Ikeo, Takenori Sasaki, Kazuyoshi Endo

## Abstract

Despite being a member of the shelled mollusks (Conchiferans), most members of extant cephalopods have lost their external biomineralized shells, except for the Nautiloids. Here, we report the result of our study to identify major Shell Matrix Proteins and their domains in the Nautiloid *Nautilus pompilius*, in order to gain a general insight into the evolution of Conchiferan Shell Matrix Proteins. In order to do so, we conducted transcriptomics of the mantle, and proteomics of the shell of *N. pompilius* simultaneously. Analyses of obtained data identified 61 distinct shell-specific sequences. Of the successfully annotated 27 sequences, protein domains were predicted in 19. Comparative analysis of *Nautilus* sequences with four Conchiferans for which Shell Matrix Protein data were available (the pacific oyster, the pearl oyster, the limpet, and the *Euhadra* snail) revealed that three proteins and six domains of the shell proteins are conserved in all Conchiferans. Interestingly, when the terrestrial *Euhadra* snail was excluded, another five proteins and six domains were found to be shared among the four marine Conchiferans. Phylogenetic analyses indicated that most of these proteins and domains were present in the ancestral Conchiferan, but employed in shell formation later and independently in most clades. Although further studies utilizing deeper sequencing techniques to obtain genome and full-length sequences, and functional analyses, must be done in the future, our results here provide important pieces of information for the elucidation of the evolution of Conchiferan shells at the molecular level.

## Introduction

Many metazoans have evolved various biomineralized tissues, both internally and externally (Cowen, 2009). Despite its maintenance cost, many metazoan species have opted to retain the presence of such tissues because they are deemed useful, for example, for structural and morphological support, mineral ions storage, and protection and defense from predators and environmental factors (Lowenstam, 1989; Simkiss and Wilbur, 2012). Among extant metazoans, two phyla have anciently evolved and are still retaining their external biomineralized shells: the mollusks (Mollusca) and the brachiopods (Brachiopoda) (Cowen, 2009). Most members of these calcifying organisms live in the marine environment, where calcium and carbonate ions are easily available as sources of the mineralized tissues (Shimizu et al, 2019).

With ca. 85000 extant members, the phylum Mollusca is one of the most successful metazoan groups. Recent phylogenomics studies have shown that a monophyletic Mollusca is comprised of two groups, the non-shell forming Aculifera (Polyplacophorans and Aplacophorans) and the external shell-forming Conchifera, which is comprised of five families grouped further into two monophyletic clades: Monoplacophorans + Cephalopods clade and Scaphopods + Gastropods + Bivalves clade (Kocot et al., 2011; Smith et al., 2011; but see Kocot, 2013 and Kocot et al., 2020). Conchiferans’ evolutionary success could probably be attributed to their ability to form mineralized external shells, which they might have acquired very early in their evolution during the Cambrian (Jackson et al., 2010; Shi et al., 2013).

The Conchiferan shell is arguably the most well studied external biomineralized structure (Marin et al., 2012). Mineralogy and microstructure studies have revealed that Conchiferan shells are mainly based on calcium carbonate, and composed of multiple calcified layers (such as the prismatic and nacreous layers) and one organic layer (the periostracum). The mechanism of shell formation is also similar among the Conchiferans: mantle tissue secretes various proteins related to mineral depositions, crystal formation breakage, pigmentation, etc. (Marin et al., 2012). Meanwhile, recent development of genomics, transcriptomics, proteomics, and other “-omics” approaches have allowed for detailed molecular characterizations of shell formation and biomineralization processes. Integration of transcriptomics or Expressed Sequence Tag (EST) analysis with proteomics have revealed a list of genes involved in biomineralization processes in the mollusks (e.g. Zhang et al., 2012; Mann et al., 2012; Marie et al., 2012; Miyamoto et al., 2013; Zhao et al., 2018). Many of such proteins are present in trace amounts inside the shell, and thus called the Shell Matrix Proteins (SMPs). Despite their small amount, the SMPs have essential roles in shell formation and structural maintenance, such as calcium carbonate nucleation, crystal growth, and choice of calcium carbonate polymorphs (Addadi et al., 2006; Marin et al., 2008).

Among the five Conchiferan orders, the evolution of the cephalopods shell is arguably the most intriguing. While the group includes famous extinct members with univalve shells such as the ammonites and belemnites, almost all extant cephalopods internalized, reduced, or completely lost their shells (such as seen in some cuttlefishes, squids, and octopods). Only *Nautilus*, the last surviving genus of the basally diverging Nautilidae (+ 416 MYA; i.e. Silurian/Devonian boundary) still have its external calcified true shells (Kröger et al., 2011). Another member of the cephalopods, the argonauts (Octopodiformes: Argonautidae) also have an external calcified shell. However, this shell is not a true shell because it lacks true shell microstructures, brittle, and most likely acquired secondarily from a shell-less Octopodiform ancestor, during the evolution of this group (Wolfe et al., 2012).

While much research on shell biomineralization genes, proteins, and protein domains have been done, most of these investigations are still biased towards bivalves and gastropods. This has hindered the elucidations of the origin and the evolution of the SMPs, including the prediction of the ancestral Conchiferan set of core protein domains needed for shell formation. Thus, in this study, we conducted transcriptomics of the mantle tissue and proteomics of the hydrophilic proteins extracted from the shell of the basal cephalopod *Nautilus pompilius* (Fig. 1). We used the transcriptome data of the mantle tissue as reference data to annotate the proteome data and thus to identify the protein sequences specifically located in the shell (the Shell Matrix Proteins; the SMPs). Comparative analyses were then conducted among the identified *Nautilus* SMPs and the publicly available representative Conchiferan SMP data of *Crassostrea gigas, Pinctada fucata, Lottia gigantea*, and *Euhadra quaesita*, in order to identify a conserved set of domains in the Conchiferan SMPs. We also conducted a SEM electron microscopy analysis of the shell of *N. pompilius* to confirm that the shell morphology, at the microstructure level, is similar to the true shells of the Conchiferans.

**Figure 1.**
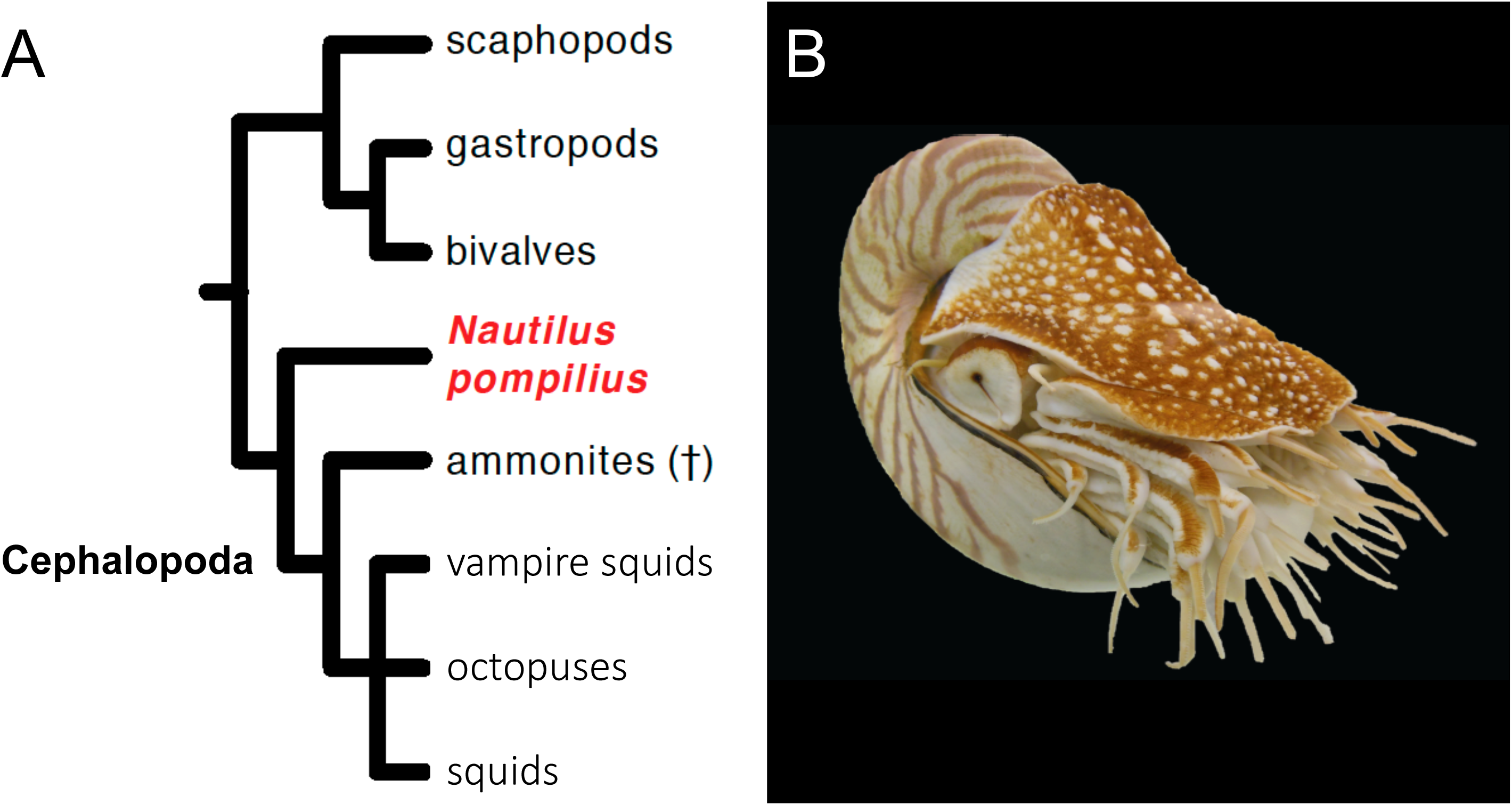
(A) Phylogeny of Conchiferans including *N. pompilius*. (B) *N. pompilius*

## Results

### The microstructure of the shell of *N. pompilius*

Our Scanning Electron Microscopy (SEM) observation confirmed that the outer shell wall of *N. pompilius* is also composed of three layers of minerals, the outer and inner prismatic layers, and the nacreous layer in between (Fig. 2A; Grégoire, 1987; Marin et al., 2012). The outer prismatic layer is the outermost layer of the *Nautilus* shell wall and comprises ~25% of the total thickness of the adult shell wall. It consists of two sub-layers, the outer sub-layer composed of small crystallite grains and the inner sub-layer composed of prism-like elongate crystals whose long axis is oriented perpendicular to the shell surface (Fig. 2B). The nacreous layer, the middle layer of the *Nautilus* shell, is the thickest layer (~70%). It is composed of numerous thin plate-like tablets, whose thickness is less than 1μm and oriented parallel to the inner shell surface. These tablets pile up one on top of another, forming columnar stacks (Fig. 2C). The inner prismatic layer comprises the innermost part of the shell. This layer is thin (~5%) and comprises prism-like elongate crystallites similar to those observed in the inner sub-layer of the outer layer (Fig. 2D).

**Figure 2.**
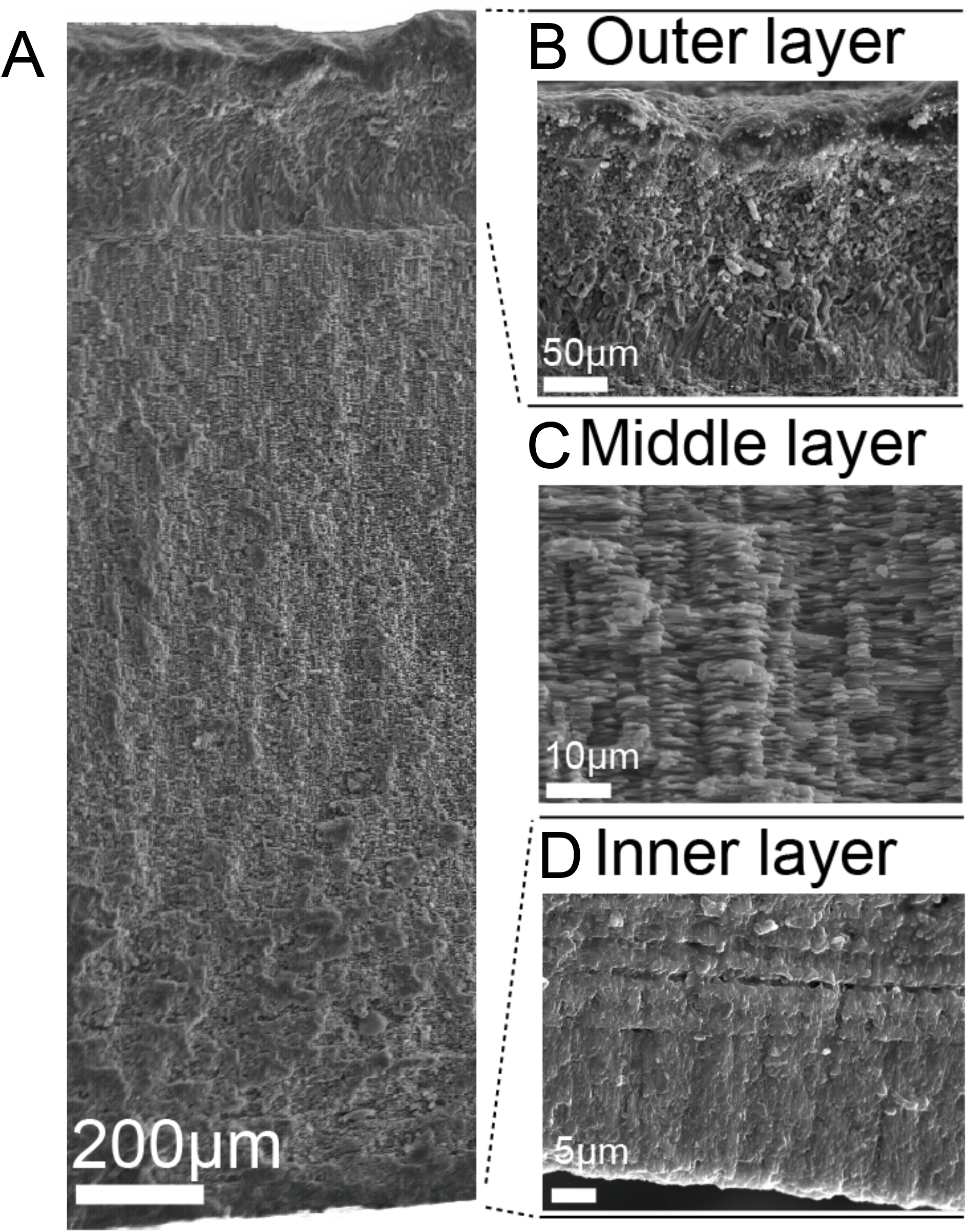
The microstructures of *N. pompilius* shell. (A) The shell microstructures of *N. pompilius*. (B) Outer prismatic layer. (C) Middle prismatic layer, (D) Inner prismatic layer.

### Transcriptomics of the mantle tissue in *N. pompilius*

We conducted transcriptome sequencing using the ION-PGM platform of seven pieces (ca. 35 mg each) of the mantle tissue in seven separated runs, resulting in about five to six million reads per run (Table 1). After sequence assembly of all reads from the seven runs combined, 48,633 contigs were obtained, with the largest contig is 13,521 bp-long, the average length of contigs 414 bp, and the N50 value 419. Of these, 11,830 contigs (24.3%) encode ORFs longer than 100aa, which 8,092 encode proteins similar to those encoded in the draft genome of O. *bimaculoides*, and 3,738 encode non-registered polypeptides/proteins, which probably include novel (previously uncharacterized) protein sequences. Five of the most abundant transcripts in the mantle tissue showed no open reading frame (ORF). Five of the most abundant transcripts with ORF were shown in Table 2.

**Table 1.** The amount and quality of the data obtained from each tissue sample

| Sample ID | individual# | tissue location | # reads | GC% |
| --- | --- | --- | --- | --- |
| NBMantle | B | mantle - random | 5,653,110 | 54.14% |
| NCMA | C | mantle - anterior | 5,958,989 | 55.43% |
| NCML | C | mantle - left | 6,434,608 | 56.06% |
| NCMR | C | mantle - right | 5,965,524 | 55.49% |
| NCMP | C | mantle - posterior | 5,299,856 | 54.85% |
| NDMP | D | mantle - posterior | 5,976,728 | 54.11% |
| NDMA | D | mantle - anterior | 6,475,168 | 55.57% |

**Table 2.** The five most abundant transcripts with ORF (in the whole mantle sample) in the mantle tissue of *Nautilus pompilius*

| contig ID | FPKM | O. bimaculoides homolog | diamond e-value | hmm domain |
| --- | --- | --- | --- | --- |
| contig_36199 | 3,124,349.10 | Ocbimv22030012m.p | 1.50E-30 | No hit |
| contig_42552 | 1,328,691.40 | None | - | No hit |
| contig_42075 | 841,970.80 | None | - | No hit |
| contig_16011 | 460,159.50 | Ocbimv22007851m.p | 2.90E-42 | No hit |
| contig_7243 | 323,500.20 | None | - | No hit |

### Sequence annotations and proteomics of Shell Matrix Proteins in *N. pompilius*

We conducted three runs of the LC-MS/MS mass spectrometer to analyze the extracted total proteins from the shell of a *Nautilus* individual for which the mantle transcriptomes were analyzed. A comparison between the obtained protein spectra from the MS/MS and the inferred protein spectra of the transcriptome contigs resulted in the identifications of 61 proteins. Of these, 14 contigs were not included in further analyses because they contain multiple translation frames, most likely frameshift error because of sequencing error.

Annotations of the remaining 47 contigs with single translation frames were conducted by doing BLASTp searches against three different databases: (1) the protein data of *Octopus bimaculoides* predicted from its genome (Albertin et al., 2015), (2) non-redundant (nr) Genbank sequence database, and (3) self-prepared database of known Shell Matrix Proteins (SMPs). The annotations were successful in identifying 27 sequences.

### Homology comparisons of the Shell Matrix Proteins among several Conchiferan mollusks

We carried out reciprocal local BLASTn searches among the Shell Matrix Proteins (SMPs) of *N. pompilius* and selected five Conchiferans for which detailed SMPs data were available (as of July 2019: the pacific oyster *Crassostrea gigas*, the pearl oyster *Pinctada fucata*, the limpet *Lottia gigantea*, and the snail *Euhadra quaesita*), in order to identify conserved proteins and conserved protein domains among the SMPs in Conchifera. The searches were conducted with the threshold of ≥50% sequence homology, and e-value of ≤e^-5^ (“Search Setting 1”). Because of the stringency of our searches, and considering our highly fragmented transcriptome sequence data, there is a possibility that we did not pick up possibly conserved protein-coding gene sequences in our data. Therefore, we also conducted reciprocal local BLASTn searches using less stringent settings following previous studies (only by setting the maximum e-value of ≤e^-5^; Shimizu et al., 2019; Zhao et al., 2018) (“Search Setting 2”).

Reciprocal local BLASTx and tBLASTn searches of the 47 SMP sequences of the *Nautilus* as queries under Search Setting 1, found 43 proteins to be specific to *Nautilus* (23 were annotated, while 20 were unknown proteins). However, the less stringent searches found 31 proteins (11 annotated, 20 unknown) to be specific to *Nautilus*. Meanwhile, searches using Search Setting 1 identified no protein, while Search Setting 2 identified additional three proteins (Pif/BMSP-like protein, CD109 antigen protein, and Tyrosinase) in all Conchiferans. Our most stringent searches identified another protein (EGF-ZP domain containing protein), and additional four (Chitinase, Peroxidase, Kunitz domain-containing protein, and *L. gigantea* LOTGIDRAFT_169029 (Chitin binding domain containing protein) by the less stringent searches, to be also shared among the four marine members, excluding *E. quaesita*. A complete list of the proteins is shown in Table 3. Meanwhile, results of the reciprocal local BLAST searches were shown as Circos charts, as shown in Fig. 3A and Supplementary Table 1 (for Search Setting 2), and Supplementary Fig. 1 and Supplementary Table 2 (for Search Setting 1).

**Figure 3.**
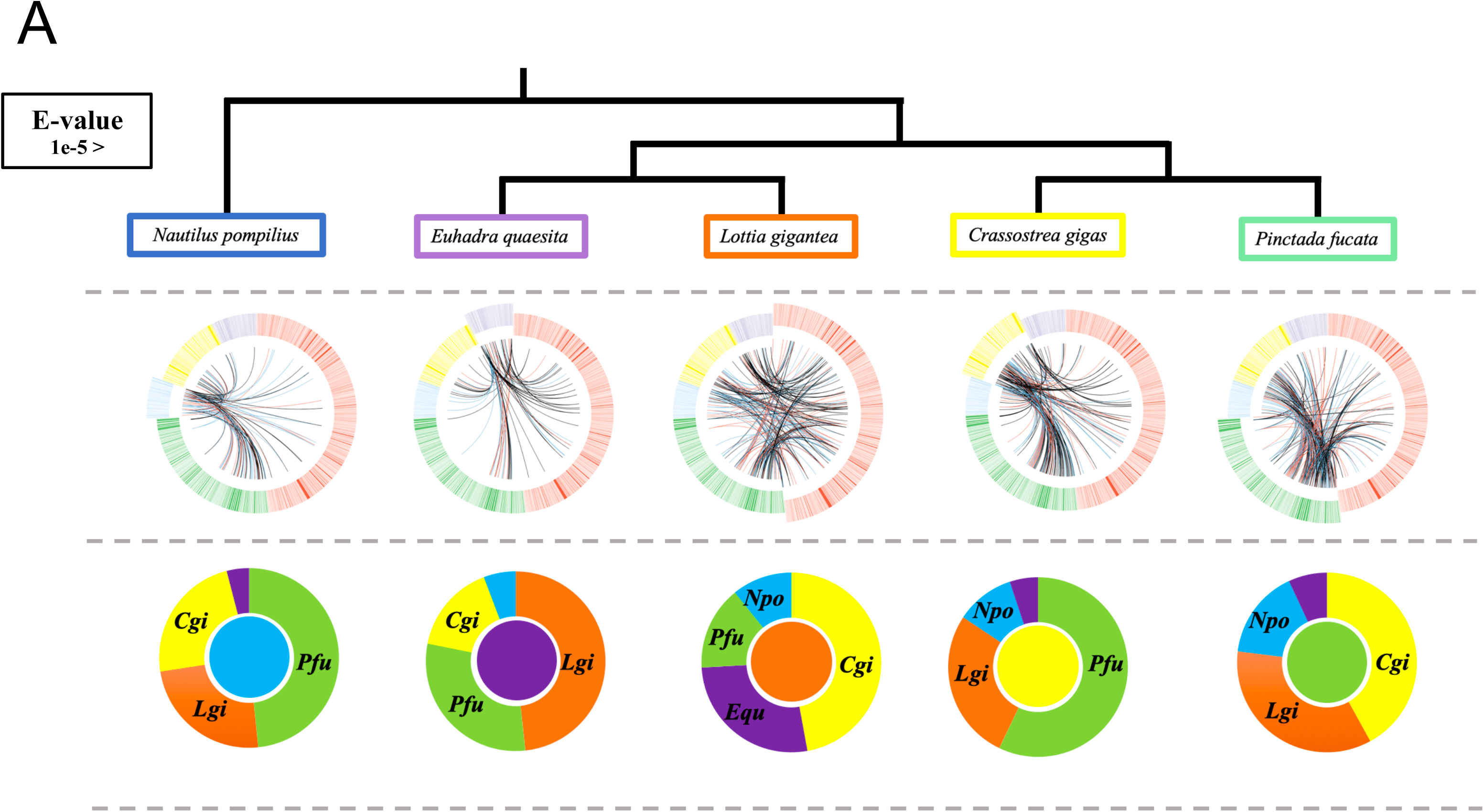
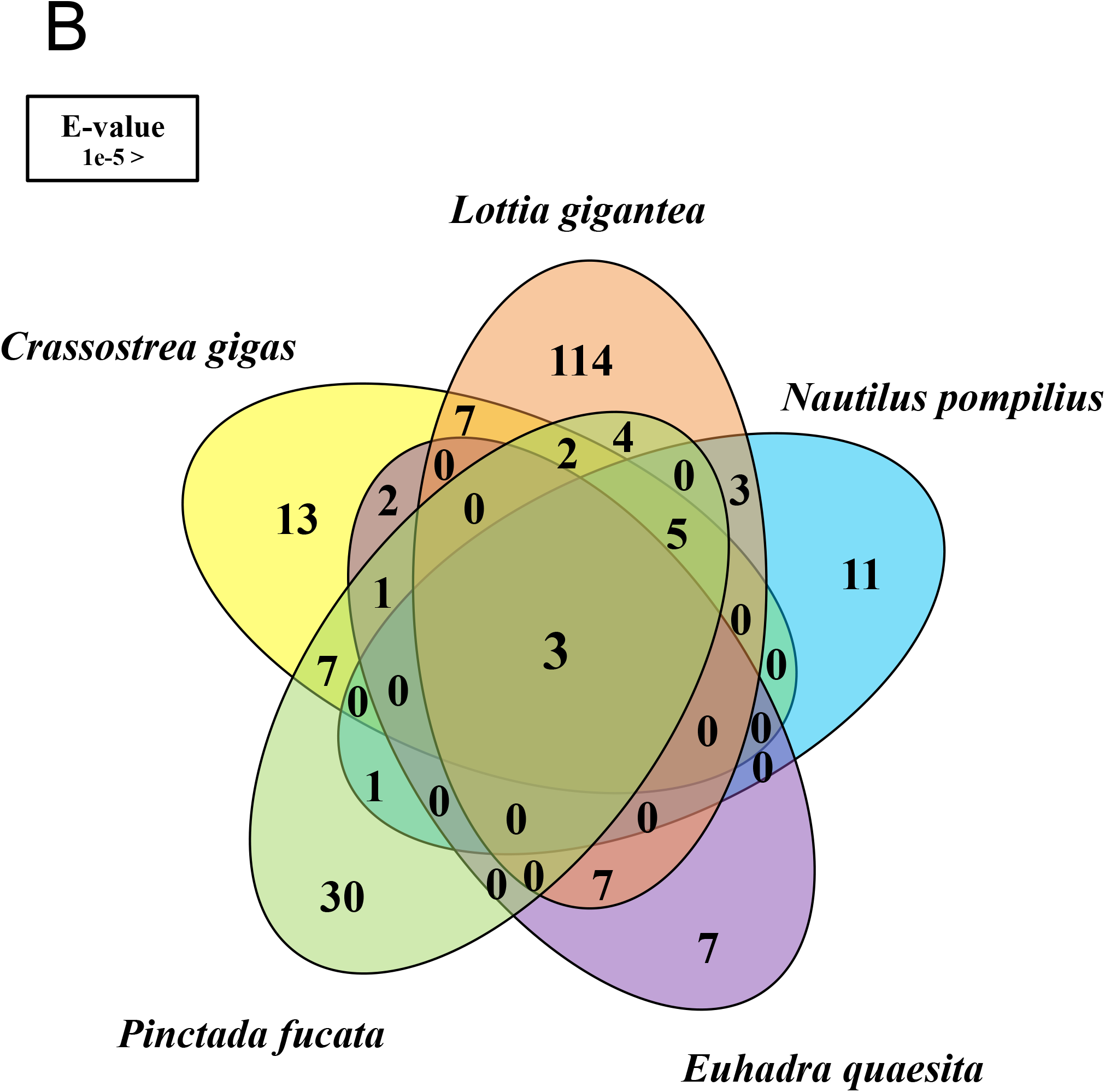
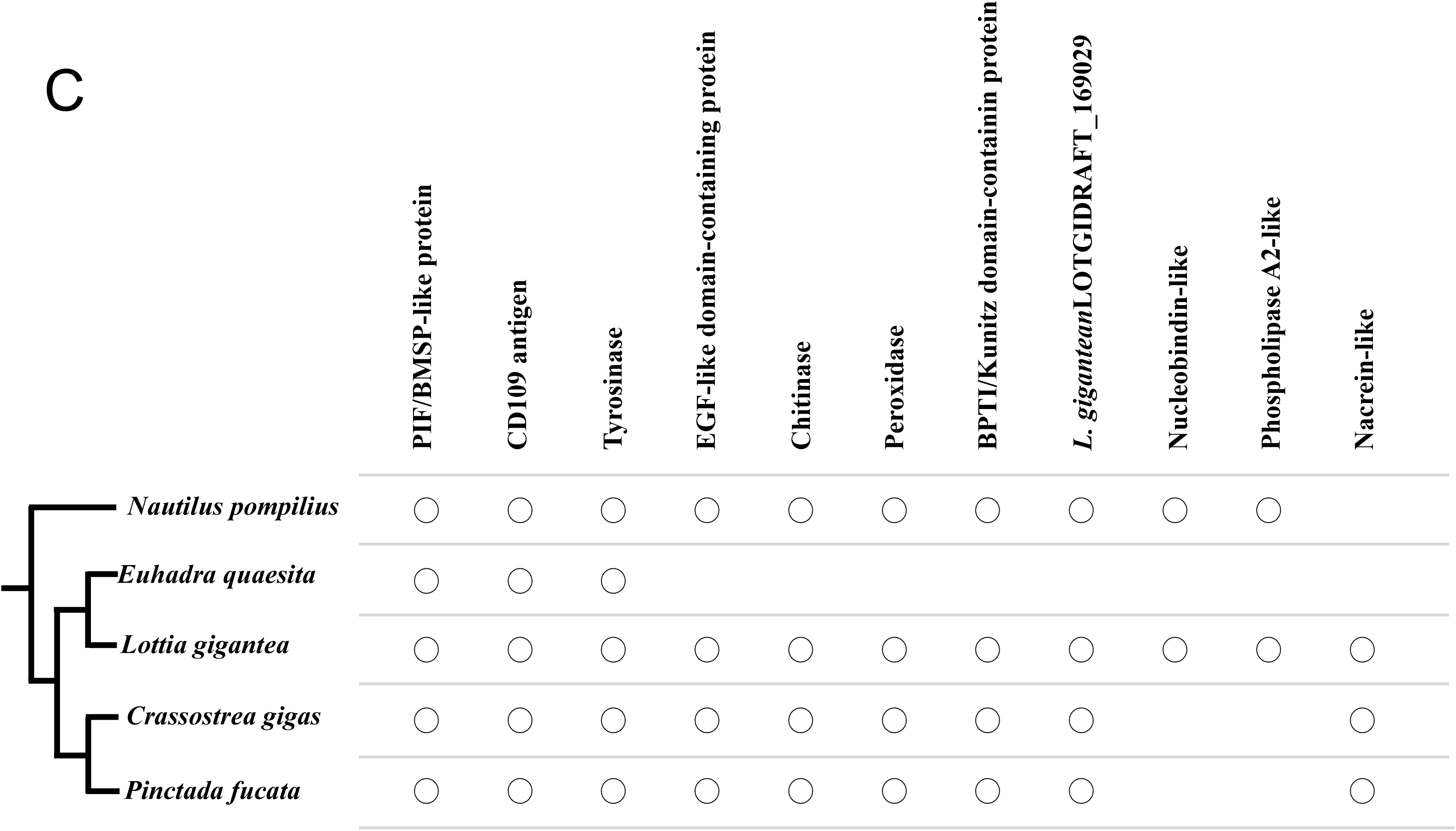
Comparisons of the Shell Matrix Proteins in several Conchiferans for which the data are available using Search Settings 1. Detailed explanation of the settings is written in the main text. (A) Schematic presentation of the homologous relationships of the Shell Matrix Proteins among five Conchiferans (*Pinctada fucata, Crassostrea gigas, Lottia gigantea*, and *Euhadra quaesita*). (B) Venn diagram showing the numbers of shared proteins identified through local BLASTp searches among the five Conchiferans. (C) Homologous proteins of the five Conchiferans compared, plotted on to the phylogeny of the animals.

**Table 3.**
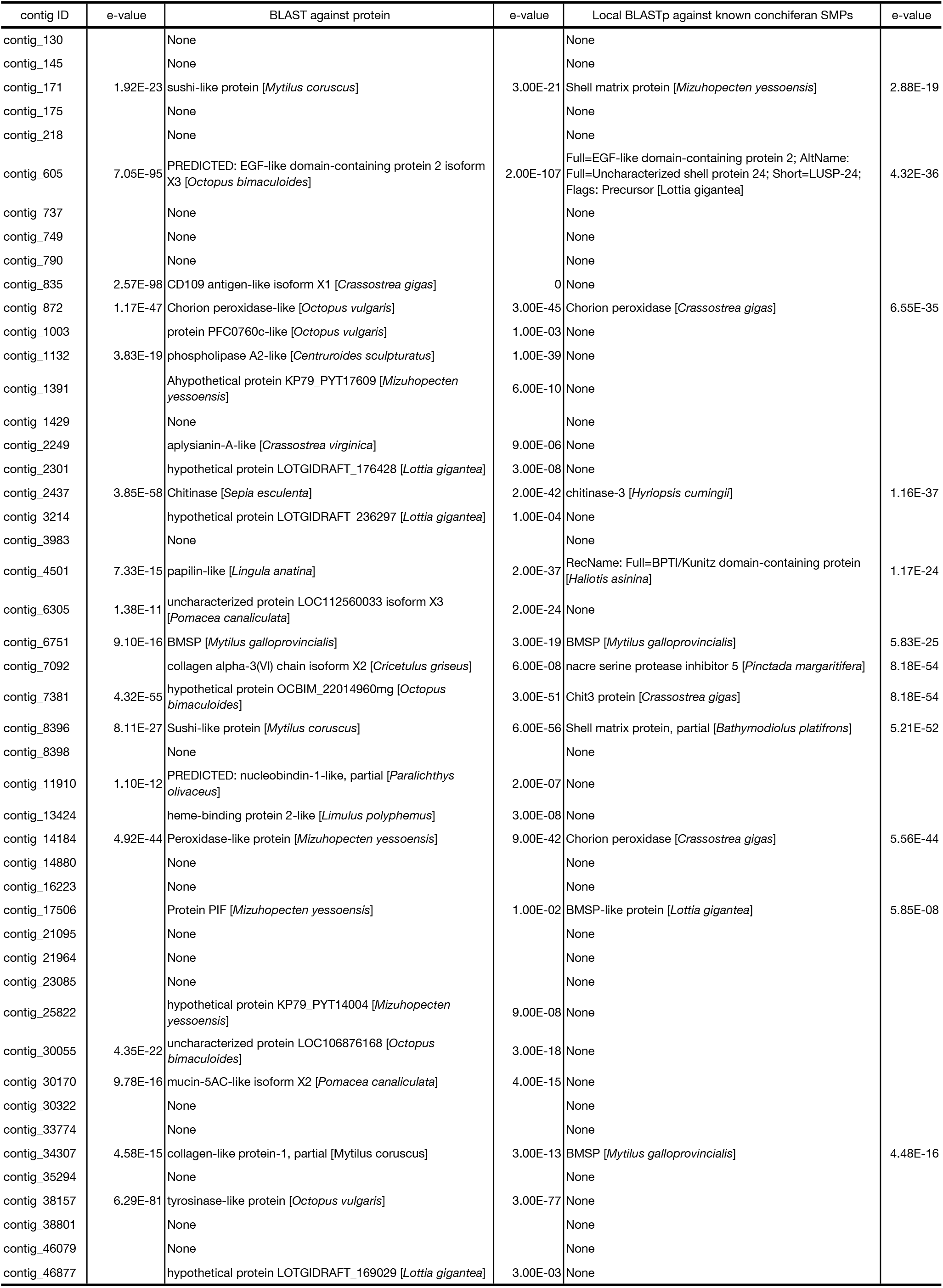
Annotation results of the 47 transcriptome contigs, which were identified as shell matrix protein-coding genes by proteome analysis

### Conserved domains of the Shell Matrix Proteins in Conchifera

Domain searches using Normal SMART (Letunic, 2018), PROSITE (Hulo et al., 2006), InterProScan (Jones *et al*., 2014), and NCBI (Altschul, 1990) databases predicted the presence of domain in 22 of the 27 annotated sequences. Meanwhile, of the unannotationable 20 contigs, domains were predicted in one contig. The diagrams showing the domains of the 22 + 1 sequences of *N. pompilius* are shown in Fig. 4A and listed in Supplementary Table 3. We manually searched for the presence of the identified domains in the other four Conchiferan Shell Matrix Protein (SMP) datasets. The result was summarized and shown in Fig. 4B, and Supplementary Tables 4–6. We found that six domains (A2M_comp, A2M_recep, Chitin-Binding Type 2 (ChtBD2), Signal peptide, Tyrosinase, and Von Willebrand factor type A (VWA)) were present in the five Conchiferans we analyzed in this study. When the terrestrial gastropod *E. quaesita* was excluded, additional six domains (An_peroxidase, Glyco_18 domain, Zona pellucida (ZP), Epidermal growth factor-like (EGF), BPTI/Kunitz family of serine protease inhibitors (KU), and Thiol-Ester bond-forming region (Thiol-ester_cl)) were found to be also shared among the four marine Conchiferans (Fig. 4B).

**Figure 4.**
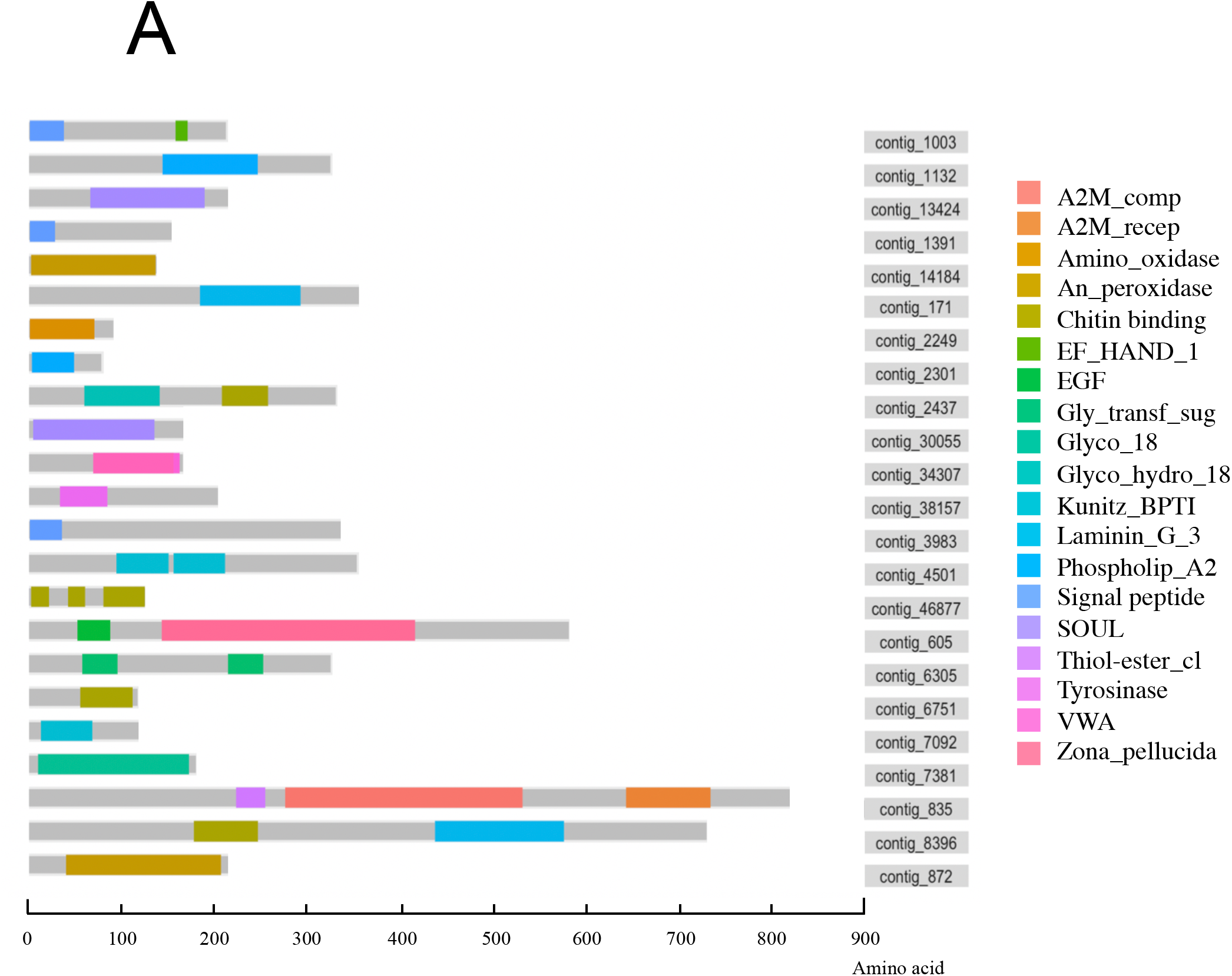
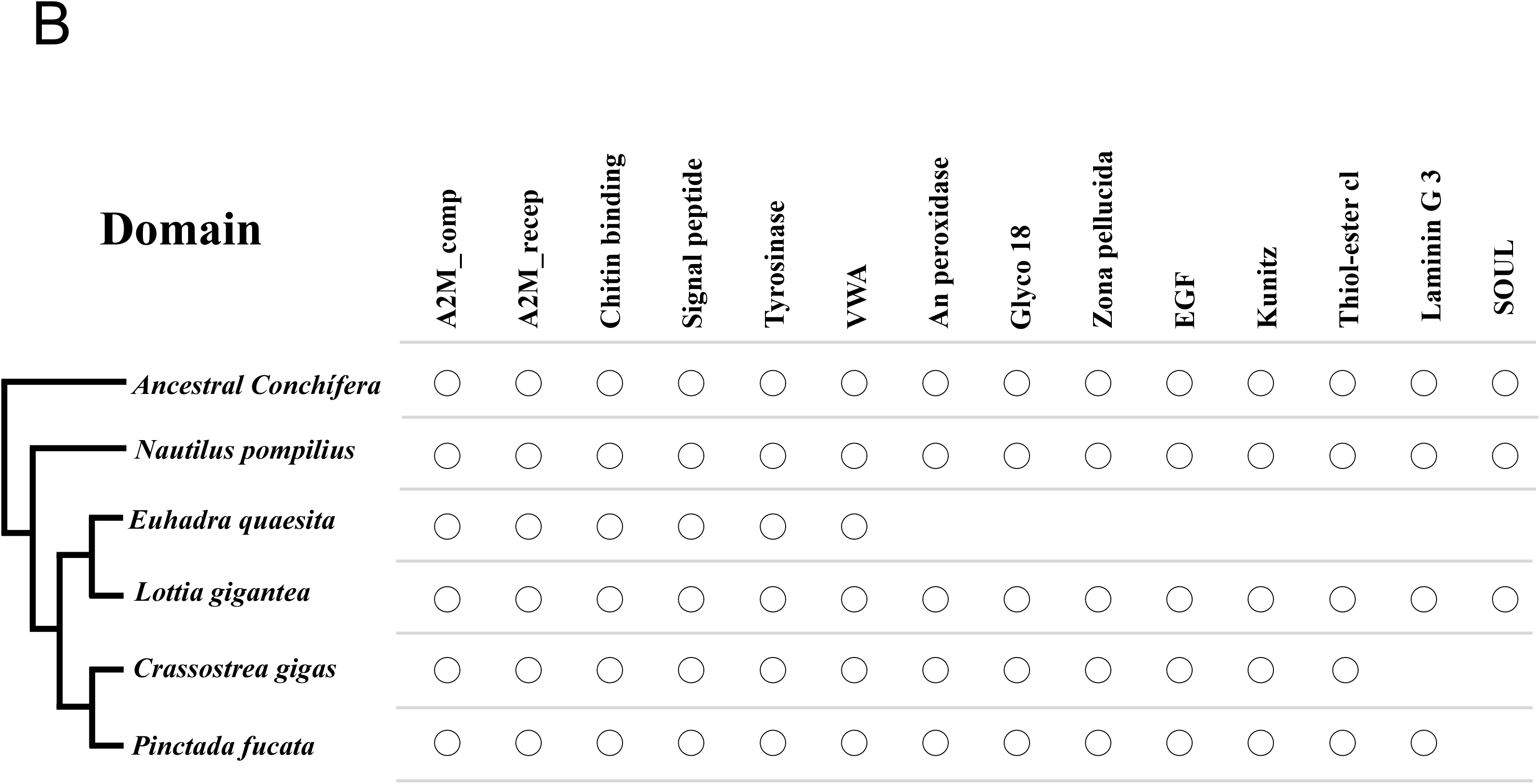
Comparisons of the domains contained in the Shell Matrix Proteins of several Conchiferans for which the data are available. (A) Schematic representations of the domains in the Shell Matrix Proteins of *N. pompilius*. (B) Shared domains in the Shell Matrix Proteins of the five Conchiferans (*Pinctada fucata, Crassostrea gigas, Lottia gigantea, and Euhadra quaesita*) compared, plotted on to the phylogeny of the animals. The reconstructed Ancestral Conchiferans most likely had all of the shared domains.

### Phylogenetic analysis of the Shell Matrix Proteins in Conchifera

As mentioned previously, we identified a total of eight proteins (Pif/BMSP-like protein, CD109 antigen protein, Tyrosinase, Chitinase, Peroxidase, Kunitz domaincontaining protein, L. *gigantea* LOTGIDRAFT_169029, and EGF-like domain containing protein) to be conserved among the four marine Conchiferans analyzed in this study. We conducted Maximum Likelihood phylogenetic inferences for the six successfully annotated proteins, in order to delve into their molecular evolutionary history. For the analyses, homologous metazoan protein sequences were mined from GenBank and UniProt, and included in the analyses. Phylogenetic analyses were conducted on the amino acid sequences of the proteins. The phylogenetic trees are shown in Fig. 5 (Pif/BMSP-like protein: Fig. 5A; CD109 antigen protein: Fig. 5B; Tyrosinase: Fig. 5C; Chitinase: Fig. 5D: Peroxidase: Fig. 5E; EGF-like domain containing protein: Fig. 5F)

**Figure 5.**
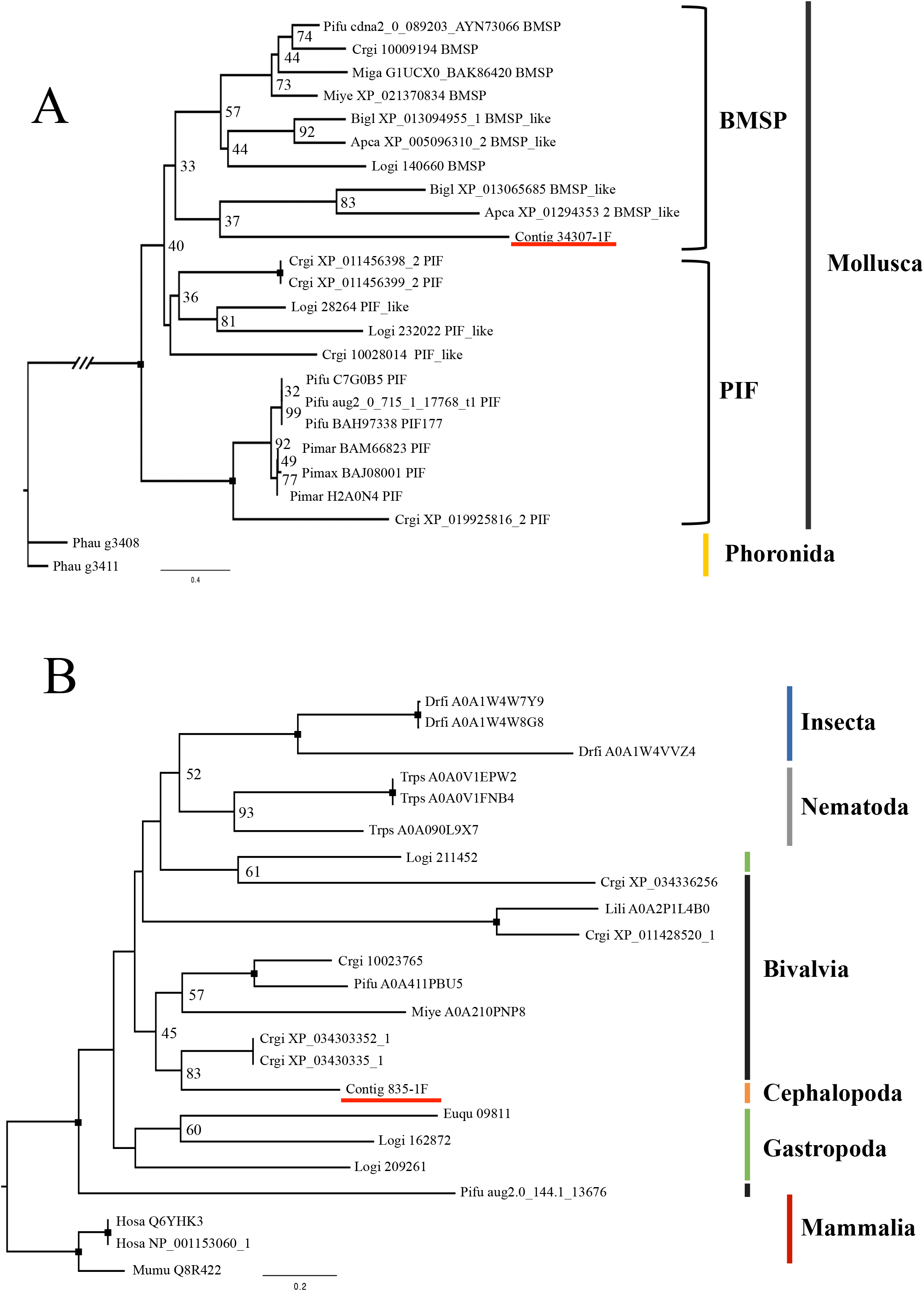
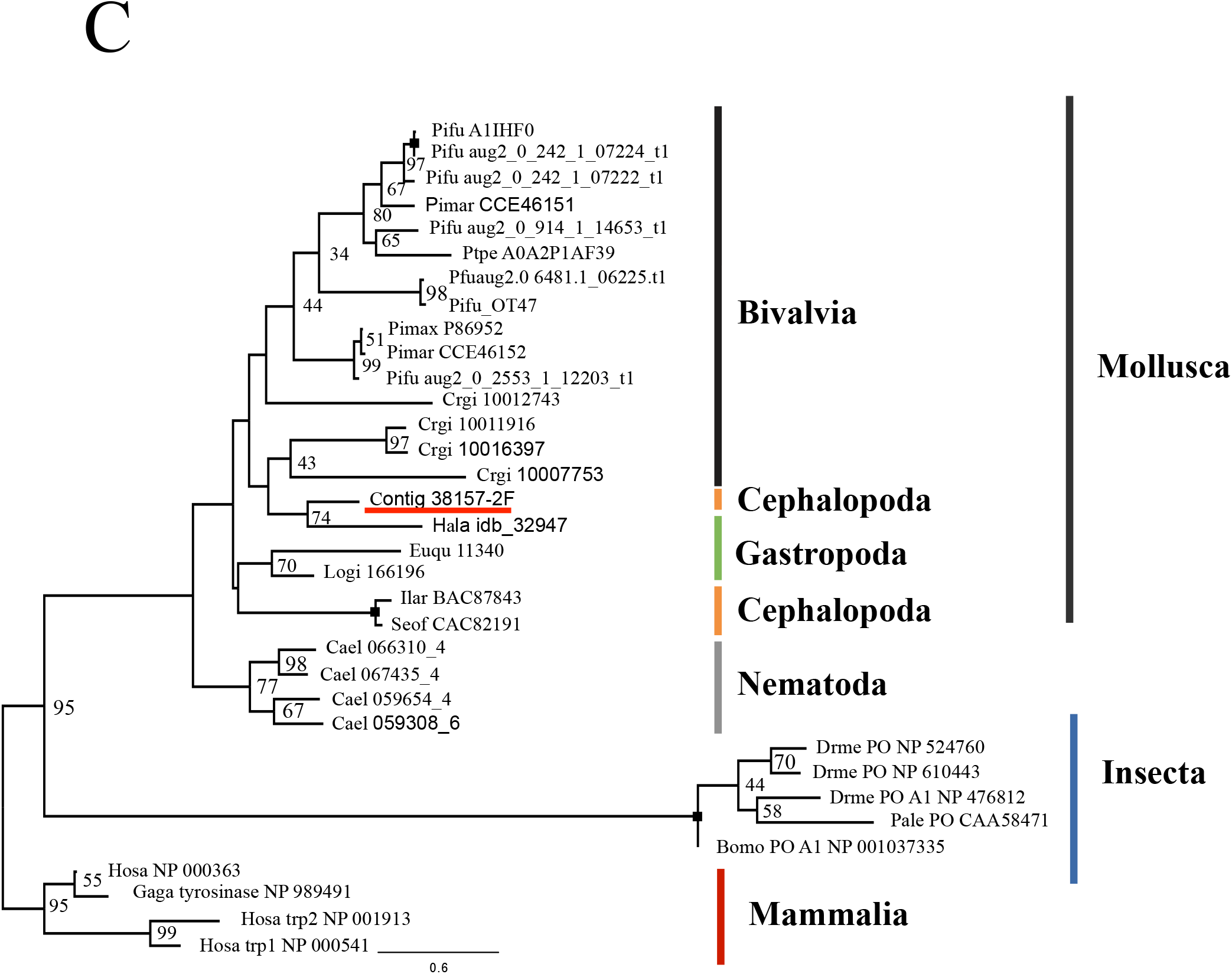
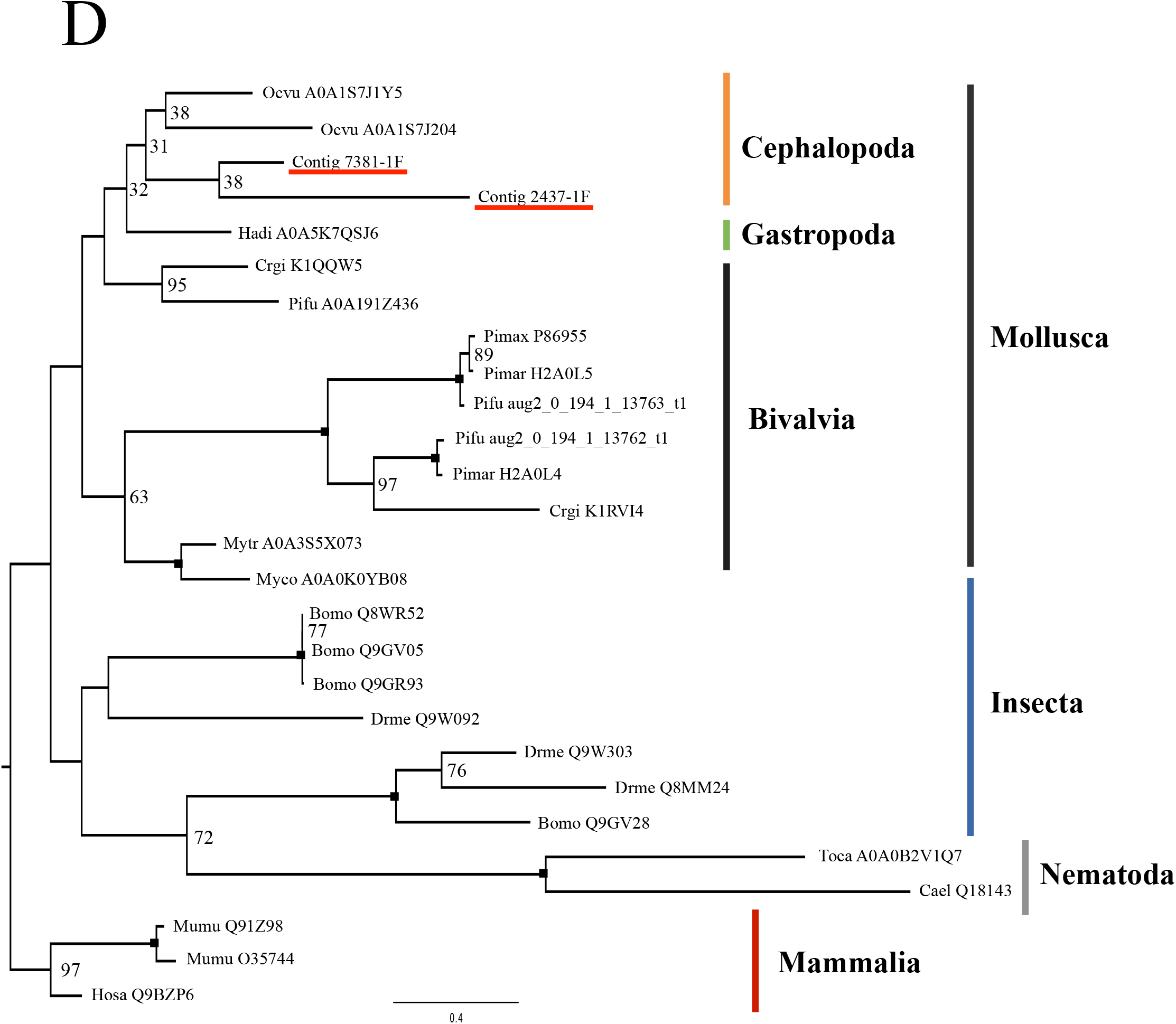
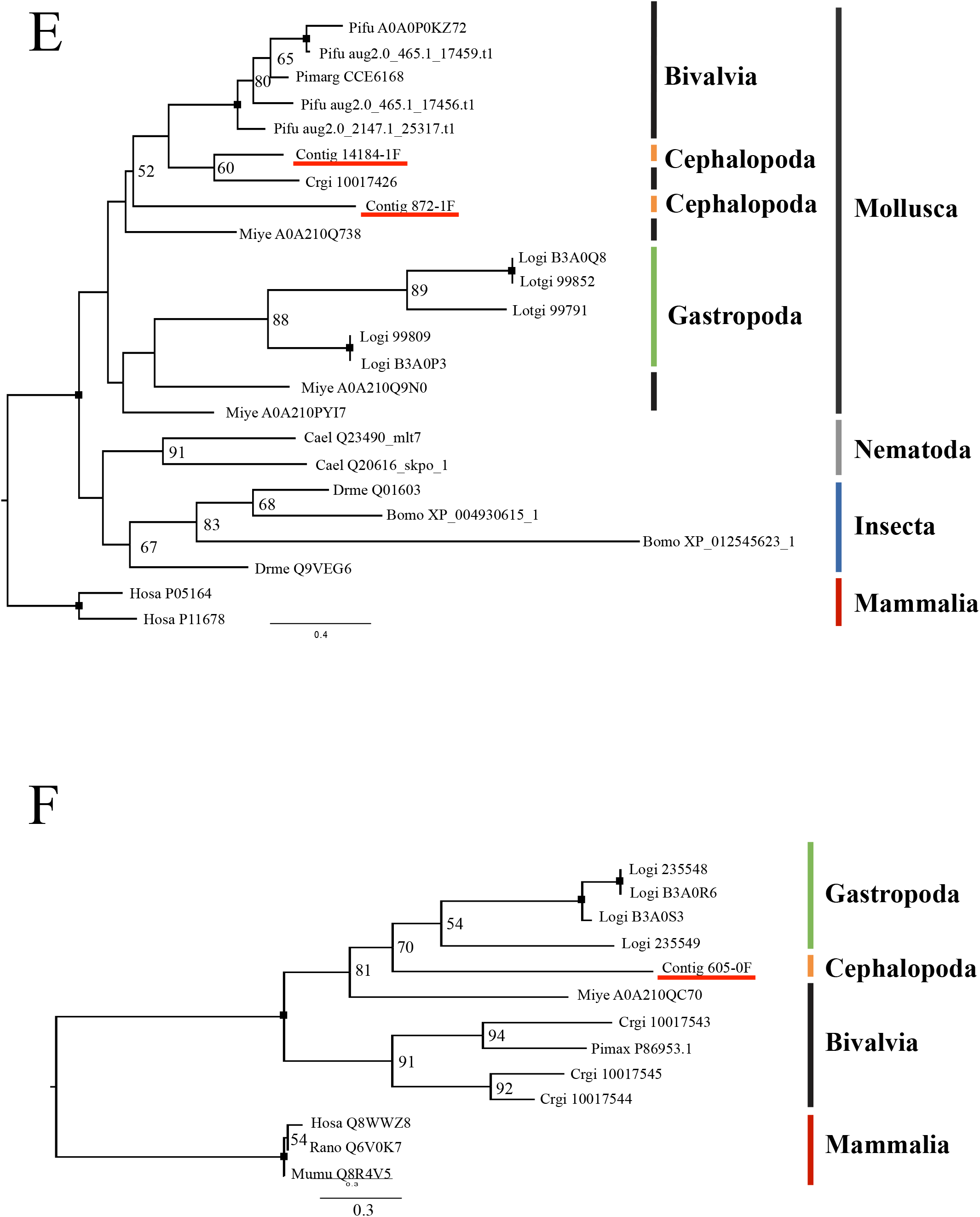
Phylogenetic trees of selected Shell Matrix Proteins. (A) The maximum likelihood tree of the Pif/BMSP amino acid sequences, inferred using the LG + Γ model with 1000 bootstrap replicates. (B) The maximum likelihood phylogenetic tree of A2M related CD109 antigen Protein, inferred using the LG + Γ model with 1000 bootstrap replicates. (C) The maximum likelihood phylogenetic tree of Tyrosinase inferred under the LG + Γ + I model with 1000 bootstrap replicates. (D) The maximum likelihood phylogenetic tree of Chitinase inferred under the LG + Γ model with 1000 bootstrap replicates. (E) The maximum likelihood tree inferred from Tyrosinase amino acid sequences under the LG + Γ model with 1000 bootstrap replicates. (D) The phylogenetic tree inferred from Peroxidase amino acid sequences under the LG + Γ model with 1000 bootstrap replicates. (F) The phylogenetic tree of the EGF-ZP Protein under the WAG + Γ model with 1000 bootstrap replicates. Bootstraps values <30% are not shown, and a black square on a node indicate 100% bootstrap support. **Abbreviations**: Pifu: *Pinctada fucata*, Crgi: *Crassostrea gigas*, Apca: *Aplysia californica*, Bigl: *Biomphalaria glabrata*, Logi: *Lottia gigantea*, Miye: *Mizuhopecten yessoensis*, Miga: *Mytilus galloprovincialis*, Phau: *Phoronis australis*, Euqu: *Euhadra quaesita*, Drfi: *Drosophila ficusphila*, Trps: *Trichinella pseudospiralis*, Hosa: *Homo sapiens*, Lili: *Littorina littorea*, Mumu: *Mus musculus*, Pimar: *Pinctada margaritifera*, Pimax: *Pinctada maxima*, Ptpe: *Pteria penguin*, Hala: *Haliotis laevigata*, Ilar: *Illex argentines*, Seof: *Sepia officinalis*, Cael: *Caenorhabditis elegans*, Drme: *Drosophila melanogaster*, Pale: *Pacifastacus leniusculus*, Bomo: *Bombyx mori*, Gaga: *Gallus gallus*, Hadi: *Haliotis discus*, Myco: *Mytilus coruscus*, Mytr: *Mytilus trossulus*, Ocvu: *Octopus vulgaris*, Toca: *Toxocara canis*, Pimarg: *Pinctada margaritifera*, Rano: *Rattus norvegicus*. Contig denotes the *N. pompilius* sequence obtained in this study.

Relatively robust phylogenetic trees were obtained for all six proteins, with most nodes supported moderately to strongly. Deeper nodes were unsupported, despite their general agreement with the accepted metazoan taxonomic classifications. The sequences form monophyletic groups at the phylum level (e.g. Mollusca), but not so at the lower taxonomic levels. All trees showed that the Shell Matrix Proteins (SMPs) are not monophyletic, and grouped together with non-SMP homologs in their consecutive phyla (Fig. 5).

## Discussion

### The shell of *N. pompilius* is a typical Conchiferan shell

Similar to other Conchiferans, the outer shells of Cephalopods are thought to also function by protecting their soft parts against predators. Shell morphological studies have indicated that outer shell breakages caused by fatal and nonfatal predatory attacks were often found in various extant *Nautilus* (e.g., Tanabe, 1988) and extinct, shelled cephalopod fossils (e.g., Takeda and Tanabe, 2015; Takeda et al., 2016). Moreover, members of Cephalopods had developed swimming ability, which had assisted their radiation both horizontally and vertically in the ocean habitat, in contrast to the rest of the marine Mollusks, which are mostly benthic. Among shelled Cephalopods, such swimming ability was acquired by the formation of chambered shells (outer shell wall + internal septa), which functioned as a hydrostatic apparatus and unique to cephalopoda (e.g., Denton and Gilpin-Brown, 1966).

The microstructures of Conchiferan shell have been classified in several ways, based on their crystalized mineral morphology and architecture (Carter, 1990). The differing classification methods however agreed on the presence of the prismatic and nacreous layers, which have been observed in the shell of all Conchiferans including *N. pompilius*, various Bivalves (e.g. Pterioidea, Mytiloidea, and Nuculoidea) and Gastropods (e.g., Trochoidea and Haliotoidea). The wide occurrence of these types of microstructures among the Conchiferans strongly suggests that the *Nautilus* shell retains some of the ancestral characters of the Conchiferan shell, and thus most likely, its biomineralization processes. The similarities in shell microstructures and morphology of *Nautilus* and other Conchiferans, and some of their functions, thus underline the importance of dissecting the molecular underpinnings of the biomineralization of the *Nautilus* shell, in order to understand Conchiferan shell evolution, at the molecular, functional, and ecological levels.

### Transcriptomics of the mantle tissue in *N. pompilius* using ION Torrent PGM is arguably enough to reveal the presence of several core Shell Matrix Proteins

In this study, we analyzed the transcriptome of several pieces of the mantle tissue obtained from three *N. pompilius* individuals. For the downstream analyses, we used a dataset built by combining all sequence reads from the seven pieces, and assembled them altogether. When analyzed together with the shell proteome data, we successfully identified 61 Shell Matrix Protein (SMP) sequences (47 SMPs = without frameshift errors), although not all of them were usable in further downstream analyses due to sequencing errors. However, the number of the obtained proteins is reasonable, when compared with other previous studies (e.g. *Euhadra quaesita* = 55, Shimizu et al., 2019; *Pinctada margaritifera* = 45, Marie et al., 2012; *Pinctada fucata* = 75, Liu et al., 2015; *Cepaea nemoralis* = 59, Mann and Jackson, 2014). One of the possible advantages of using a shallow system for transcriptome sequencing is that, most of the sequences we obtained here were probably the most abundantly expressed transcripts, and thus, major SMPs, and not background expression genes accidentally picked-up. However, using a shallow next generation sequencing system such as ION-PGM also brings some disadvantages. For example, failure in domain predictions and annotations of several SMP contigs were probably because they were too fragmented and thus the sequences were incomplete, causing the annotation programs to be unable to detect any active domain sequences. There is also a possibility that sequencing errors might have caused mis-*in silico*-translations of some contigs. Of course, however, the possibility that some of the contained domains were unpredictable because they were novel domains, and that the 13 protein sequences are novel, previously uncharacterized proteins, cannot be eliminated by our present results.

For example, in this study, we were also unable to identify the only previously reported SMPs of the *Nautilus* thus far: Nautilin-63, which was extracted from the hydrophilic fraction of the shell of a congener of *N. pompilius, N. macromphalus* (Marie et al., 2011). This is probably caused by the shallowness of the sequencing system we presently employed. However, the possibility that this protein is species specific cannot be denied. Future analyses are still needed to see if Nautilin-63 is a major protein in all Nautiloids, or specific to *N. macromphalus*.

Therefore, in order to obtain the complete picture of SMPs in *N. pompilius*, further studies using deep transcriptome sequencing systems such as Illumina, and proteomics analyses of both the hydrophilic and hydrophobic component of the SMPs, are still needed in the future.

### Homology comparisons and the evolution of the Shell Matrix Proteins among several Conchiferan mollusks

Homology searches among several Conchiferan mollusks for which the Shell Matrix Proteins (SMPs) have been studied as of July 2019 (the pacific oyster *Crassostrea gigas*, the pearl oyster *Pinctada fucata*, the limpet *Lottia gigantea*, and the snail *Euhadra quaesita*) revealed that three proteins (Pif/BMSP-like protein, CD109 antigen protein, and Tyrosinase; Fig. 3C) shared among the Conchiferans. The three proteins are known to be very important in maintaining shell structures. For example, the Pif/BMSP proteins are involved in the formation of the nacreous layer of the shell, and thus crucial in forming and maintaining shell structure (Miyamoto et al., 2013; Suzuki et al., 2009; Suzuki et al., 2011). Pif and BMSP are composed of signal peptide, von Willebrand factor Type A domain (VWA), and Chitin-binding domains. VWA domain has function of the protein-protein interaction, Chitin-binding domain has the interaction with calcium ions in calcium carbonate (Suzuki et al., 2011). Meanwhile, Tyrosinase (both as a protein and a domain) is known to be involved in pigmentation (Nagai et al., 2007; Yao et al., 2020), and found in all mollusks compared in this study. Tyrosinase involvement in pigmentation is not only in the shell, but the protein was probably recruited and included inside the shell matrices to form the diverse coloration and patterns of the shell. In mammals including humans, the CD109 antigen protein is known to be involved in mineralized tissue formation, by being involved in osteoclast formations (Wang et al., 2013). Molecularly, it is a protease inhibitor, and it works by regulating TGF-beta receptor expression, TGF-beta signaling and STAT3 activation to inhibit TGF-beta signaling (Finnson et al., 2006; Litvinov et al., 2011).

Besides the three proteins detailed above, when the land snail *Euhadra quaesita* was excluded in the reciprocal BLASTx searches, another five proteins (EGF-ZP domain containing protein, Chitinase, Peroxidase, Kunitz domaincontaining protein, and *L. gigantea* LOTGIDRAFT_169029 (Chitin binding domain containing protein) were found to be conserved among the marine Conchiferans (Fig. 3C). While it is very enticing to suggest that the difference in the types of proteins inside the shell matrices were caused by adaptation to the terrestrial environment, our analyses reported here cannot conclusively suggest so because of the differences in sequencing methods, sequencing depths, and completeness of the data compared. However, previous reports have suggested that the proteins reported as conserved among the marine Conchiferans were also probably important during shell formation. For example, the EGF-ZP domain-containing protein, Chitinase, and Peroxidase are suggested to be involved in the formation of calcium carbonate crystals in the shell (Iwamoto et al., 2020, Kintsu et al., 2017, Liao et al., 2019, Hohagen and Jackson, 2013). Future functional studies on these proteins, including the presently unknown *L. gigantea* LOTGIDRAFT_169029, must still be conducted in the future to investigate their specific functions during Conchiferan shell formation.

Two proteins, Nucleobindin-like and Phospholipase A2-like proteins, were shown to be shared only between the limpet *L. gigantea* and *Nautilus*. Nucleobindin is known to be related to calcium ion binding in humans (Gaudet et al., 2011). Phospholipase A2 is a hydrolyzing enzyme which function of cleaving phospholipids depends on the presence of calcium ions (Dennis, 1994). While the specific function of both enzymes during shell formation and biomineralization has never been assessed, we could deduce that both enzymes are probably related to the calcification process of the shell. However, our analyses did not find these two enzymes in the shell matrices of other Conchiferans. This could be attributed not only to the exhaustiveness of data, but also to possible evolutionary scenarios, where the two genes were either lost by the other Conchiferan groups, or independently or recruited by *L. gigantea* and *Nautilus*. Interestingly, the traditional view of Molluscan taxonomy puts Gastropods as the sister group of Cephalopods (e.g. Yochelson et al., 1973, Salvini-Plawen and Steiner, 1996.). It is also to be noted that we found two Phospholipase A2-like proteins in *Nautilus*.

We did not find Nacrein-like protein in our *Nautilus* transcriptome and proteome data, although it is present in all other marine Conchiferans compared in this study. Interestingly, this protein is considered as one of the major soluble SMPs, and thus should be detected in our present data because we analyzed only the hydrophilic fraction of the *Nautilus* SMPs. However with our present data, we cannot say for certain that it is absent in the *Nautilus*. We believe that this protein should be present in all Conchiferans, although undetectable in our present *Nautilus* data. Future studies including the hydrophobic fraction of the SMPs of *Nautilus* using different sequencing platforms is still needed to clarify this issue.

Based on the information we presently obtained, we can deduce the Conchiferan core set of SMPs (Fig. 3C). However, phylogenetic analyses of the six proteins (Fig. 5A–F) showed that the SMPs were not monophyletic, as what would be expected if the proteins were specifically recruited once in the ancestral Conchiferan, to be used in shell formation. We found that the SMPs were not monophyletic even among closely related taxa/species. Therefore, with our present finding, we can deduce that the same proteins were probably recruited multiple times in various taxa across Conchiferans, from preexisting proteins, which functions and structures were probably useful and easier to tinker for the formation of biomineralized structures.

### Homology comparisons and the evolution of the Shell Matrix Proteins domains

From the 47 protein sequences we obtained from the shell of *N. pompilius*, we identified the presence of 19 domains (Fig. 4A). When compared with other the Shell Matrix Protein (SMP) data of the other Conchiferans analyzed in this study, we identified that five domains were conserved among all Conchiferans, and five additional domains were conserved among the marine species (Fig. 4B), and three domains were found only in *Nautilus*. They are common domains usually found in many proteins, including those unrelated to the biomineralization process in metazoans. However, we can deduce that the proteins containing the domains were recruited for shell formation, because the domains’ known functions are most likely related to one or several activities/events during shell formation and maintenance, including the biomineralization process.

### The Shell Matrix Proteins of *N. pompilius*

In this study, of the 47 proteins we successfully identified using both the transcriptome and proteome data, only 27 were successfully annotated. We were unable to annotate the 20 protein sequences, probably because they are too short, or previously uncharacterized novel protein sequences. However, the lack of sequence information thus prohibits us to deduce if the sequences were unique to *Nautilus*, or shared with other organisms we compared in this study.

Meanwhile, of the 27 sequences we annotated, we found 11 proteins (PFC0760c-like protein [*Octopus vulgaris*], Phospholipase A2-like [*Centruroides sculpturatus*], heme-binding protein 2-like [*Limulus polyphemus*], hypothetical protein KP79_PYT17609 [*Mizuhopecten yessoensis*], uncharacterized protein LOC110465975 [*Mizuhopecten yessoensis*], hypothetical protein KP79_PYT14004 [*Mizuhopecten yessoensis*], mucin-5AC-like isoform X2 [*Pomacea canaliculata*], uncharacterized protein LOC112572957 [*Pomacea canaliculata*], uncharacterized protein LOC112560033 isoform X3 [*Pomacea canaliculata*], and two Sushi-like protein [*Mytilus coruscus*]) to be present only in the shell matrix of *N. pompilius* (Table 3). With only our present data, we are unable to actually say if the lack of these proteins in other Conchiferans biological or technical. For example, it is possible that the protein shared between *Nautilus* and the octopus (hypothetical protein OCBIM_22021924mg [*Octopus bimaculoides*]) are actually a protein sequence specific to the Cephalopods, while the heme-binding protein 2-like [*Limulus polyphemus*] are shared between Cephalopods and the Limulid Arthropods, the horseshoe crabs. Comprehensive future studies involving molecular evolution studies, comparative genomics, and functional analyses comparing these proteins are needed in order to obtain conclusive insights regarding their functions, and their specificity (or non-specificity) in the *Nautilus*.

It is also to be noted that we also found the EGF and ZP domains-containing protein in *N. pompilius* (Fig. 4A). The presence of the homologs of this protein in all Conchiferan SMPs including the basal cephalopod *Nautilus* might have underlined the importance of this protein during Conchiferan shell formation (Feng et al., 2017).

## Ethical statement

All experiments were conducted in accordance to the guidelines and protocols of The University of Tokyo, in order to ensure proper and humane treatments of the experimental animals sacrificed during the course of this study.

## Funding and research support

During the course of this study, DHES were initially supported by the JSPS Foreigner Postdoctoral Fellowship (F02330) with the entailing research grant awarded to KE (12F02330), and then partially by the FY2016 Research Grant for Chemistry and Life Sciences (The Asahi Glass Foundation), and the FY2017 Research Grant for Zoology (Fujiwara Natural History Research Foundation), both awarded to DHES. The study was also supported partially by the FY2018 Grant-in-Aid for Scientific Research (C) (Grant number: 18K06363) awarded to MAY (PI) and DHES (Co-PI). KE was partially supported by FY2011 Grant-in-Aid for Scientific Research (A) (Grant number: 23244101) and FY2011 Grant-in-Aid for Challenging (Exploratory) Research (Grant number: 23654177). KS was supported by Grant-in-Aid for JSPS Fellows (Grant number: 12J09867).

DHES would like to thank the constant support and invaluable advice given by present and former members of the Setiamarga lab at NITW, former members of Endo lab at the University of Tokyo, and Takeshi Takeuchi (OIST).

## Author contributions

DHES conceived the idea, initiated, and together with KE, managed the course of the study. DHES, HK, MAY, and KI conducted data analyses. DHES, TS, KS, and YT euthanized and dissected samples, which were then vouchered by TS at the museum. YT conducted shell microstructure analyses. DHES, HK, KS, YI, and KK conducted molecular works. DHES and HK wrote the first draft of the manuscript, which were then edited further by DHES, HK, MAY, YT, and KE. All authors confirmed the content of the final version of this manuscript.

## Materials and Methods

### Microstructure observations of the shell of *N. pompilius*

The microstructure of the outer shell of *N. pompilius* was examined by SEM (VE-8800, Keyence, Osaka, Japan). Samples of ~1 cm2 were removed from the individual shell and their fracture surfaces were examined. Prior to the SEM observation, they were treated in etching with hydrochloric acid for 30 seconds. All samples were coated with platinum.

### Sample collection and RNA extraction

We obtained three individuals of *N. pompilius* from a local dealer for aquarium shops in Japan. The samples were obtained from The Philippines. We obtained these samples at the end of 2011 and beginning of 2012, before the inclusion of this species in the CITES list and thus prior to the protected status of this species under the Washington agreement. First, we sedated the individuals in 2% ethanol in cold sea water for ca. 10 minutes (Butler-Struben et al., 2018). Afterward, we removed the shells of the individuals, and cut out pieces of the mantle tissue (ca. 25–35 mg each; Table 1) on ice, and stored them in ISOGEN (Nippon Gene Co. Ltd., Tokyo, Japan) at −80°C. Total RNA was extracted from the tissue samples using ISOGEN and the RNeasy kit (Qiagen), and was stored in −80°C until further transcriptome analyses. The rest of the body of the individuals were euthanized by freezing them in −80°C, and then preserved in formalin, to be later stored as vouchered specimens at The University Museum, The University of Tokyo, Japan.

### Transcriptome analyses

Transcriptome sequencing of the mRNA extracted from the seven tissue samples, using the Ion Torrent PGM platform (Thermo Fisher Scientific) was outsourced to the Center for Omics and Bioinformatics, The University of Tokyo. Afterward, the obtained raw reads from the seven libraries made from the seven tissue samples were combined, and then assembled using the CLC assembly cell with the default settings on the Maser computing system, Data center for cell innovation, National Institute of Genetics (Kinjo et al. 2018). The Maser analytical pipelines on the National Institute of Genetics Cell Innovation program (http://cell-innovation.nig.ac.jp/) were used for the following functional estimations of the assembled CLC contigs. For expression profiling, FASTQ reads were aligned to the CLC contigs using the TMAP mapping program (https://github.com/iontorrent/TS/tree/master/Analysis/TMAP). Raw read sequence data will be available in the DNA Data Bank of Japan (DDBJ).

### Proteome analyses of total hydrophilic protein from the shell of *N. pompilius*

Shell of a *Nautilus* individual, for which the mantle transcriptomes were analyzed, was first shattered into pieces using a hammer. The shell pieces were cleaned from any organic tissue by incubation in a 2M NaOH overnight, and a thorough washing with Milli-Q water 10 times. Cleaned shell pieces were then ground into powder, and then slowly decalcified using 0.5 M EDTA as the chelating agent, at 4°C for 3 days. Total hydrophilic proteins of the shell were extracted using the 3 kDa Amicon Ultra Centrifugal Filter Unit.

After digestion into short peptides by trypsin (Promega), the samples were analyzed using a DiNa nanoLC system (KYA Technologies, Tokyo, Japan) and a LTQ Orbitrap mass spectrometer (Thermo Fisher Scientific). Identifications of obtained spectra were conducted by conducting a search on a self-prepared protein sequence database using the spectra as queries, using the SEQUEST program in Proteome Discoverer version 1.2 (Thermo Fisher Scientific). The self-made protein sequence database contained bioinformatically translated sequences of the assembled transcriptome contig data from the mantle tissue. These “theoretical” protein sequences were then fragmented into peptides *in silico* to simulate digestion by trypsin, in order to obtain the theoretical mass of peptides and MS/MS spectra. Spectrum data searches matched the actual experimental data of the actually obtained LC MS/MS spectra of the Shell Matrix Protein polypeptides, with the theoretical spectra database, resulting in the identification of candidate protein sequences from the database. Only transcriptome-based protein sequences matched by at least two LC MS/MS polypeptides were selected as potential Shell Matrix Proteins. Detailed methods and parameters for analyses were described in Elias and Gygi (2007), Isowa et al. (2015), and Shimizu et al. (2019).

### Characterizations of the Shell Matrix Proteins of N. pompilius

Sequence annotation was performed by conducting BLASTp and BLASTx searches on the nr databases of Genbank and a database of published Conchiferan Shell Matrix Protein sequences, which we compiled ourselves by expanding the dataset of Arivalagan et al. (2017) and Feng et al. (2017).

Protein domains were predicted using multiple online tools: SMART (http://smart.embl-heidelberg.de/), PROSITE (https://prosite.expasy.org/), InterProScan (https://www.ebi.ac.uk/interpro/search/sequence/), and Pfam (HMMER v3.3; e-value <1.0e-5; http://hmmer.org/). Signal peptides were predicted using the online tool SignalP (Petersen et al. 2011). Predicted domains were visualized using an R script (Fig. 4A).

### Comparative analysis of Conchiferan Shell Matrix Proteins

In order to identify conserved protein sequences among the five Conchiferan species analyzed in this study, the annotated 47 Shell Matrix Protein sequences of *N. pompilius* were used as queries in reciprocal local BLASTx and tBLASTn searches, against four molluscan for which the Shell Matrix Protein sequence data are already published (71 *Crassostrea gigas* proteins (Zhao et al, 2018); 159 *Pinctada fucata* proteins (Zhao et al 2018); 311 *Lottia gigantea* proteins (Mann et al 2012); 55 *Euhadra quaesita* proteins (Shimizu et al 2019)) (e-value <1e-5 and threshold ≥50%: “Search Setting 1”, e-value <1e-5: “Search Setting 2”). The result was visualized as Circos charts using the software Circos-0.69-9 (http://circos.ca/) (Fig. 3A).

The presence of homologous domains was confirmed manually, based on our reciprocal local BLAST result. The result was summarized and presented as a Venn diagram (Fig. 4B).

### Phylogenetic analyses of the Shell Matrix Proteins

Phylogenetic analyses were conducted on a total of seven Shell Matrix Proteins obtained in this study (Tyrosinase, An-peroxidase, Chitinase, A2M receptor-domain containing Antigen-like protein, EGF-ZP, and BMSP). In order to do so, homologous amino acid sequences of each protein of various organisms were data-mined from UNIPROT (https://www.uniprot.org/), including molluscan SMPs (if available / relevant), and non-SMPs. The presence of homologous domains in the sequences was confirmed using HMMER v3.1b2 (http://hmmer.org; e-values < 1.0e-5). These sequences were then aligned using the online version of MAFFT v7.310 (http://mafft.cbrc.jp/alignment/server/index.html; Katoh et al., 2002), with the g-INS-i algorithms to allow for global alignment (Katoh et al., 2005). Sequences were edited using the online version of GBlocks v.091b (Castresana, 2001) under the least stringent settings. Model selection was conducted on MEGA v10 (Tamura et al., 2011). Maximum Likelihood trees were inferred using the GUI version of RAxML (Silvestro et al 2012), with the rapid tree search setting and 1000 bootstrap replications, using the best fitting amino acid substitution model. The selected model for each protein is written directly in the figure showing the phylogenetic tree.

## Supplementary Figure Legends

**Supplementary Figure 1.**
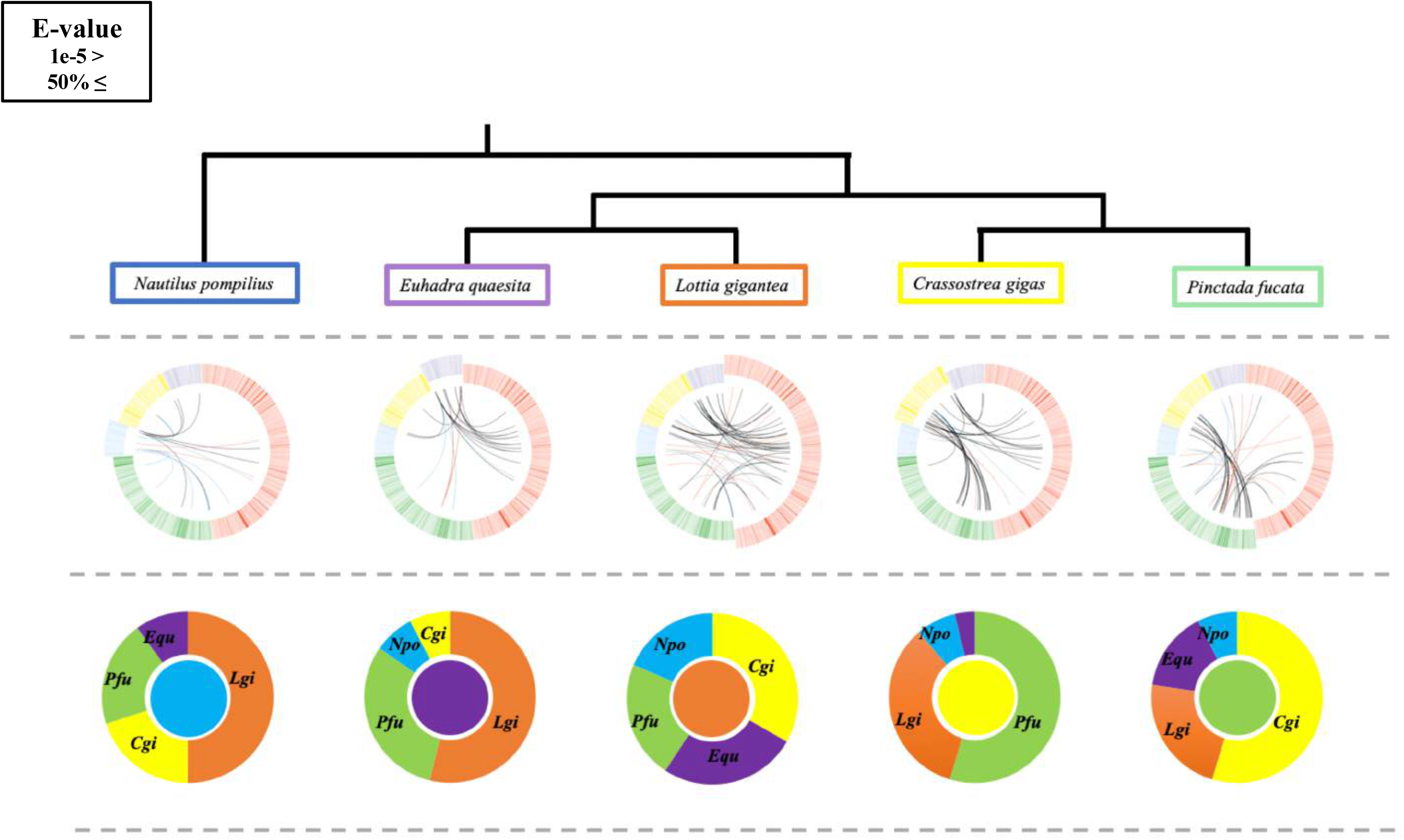
Comparisons of the Shell Matrix Proteins in several Conchiferans for which the data are available using Search Settings 2. Detailed explanation of the settings is written in the main text.

**Supplementary Table 1.**
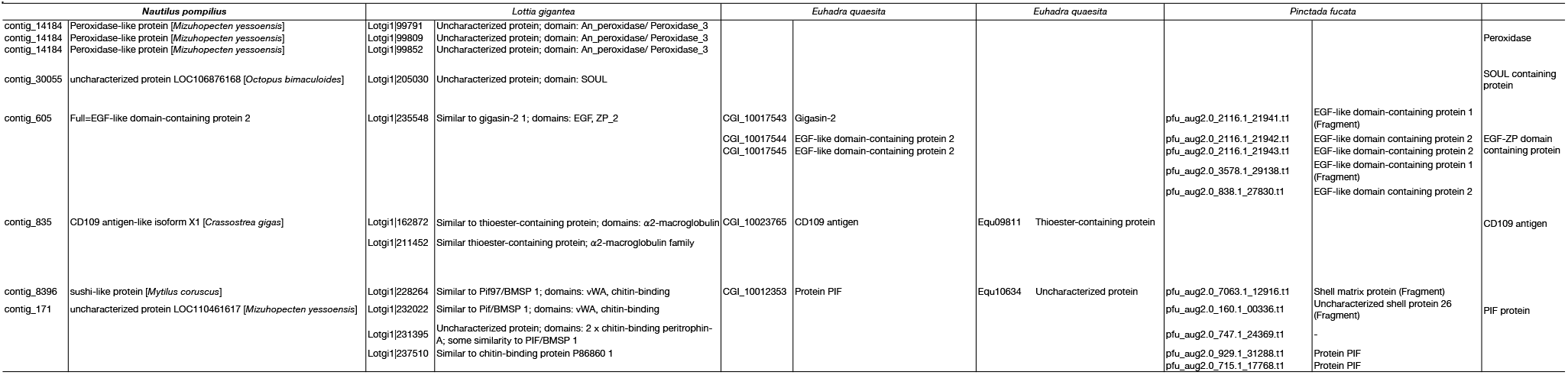
Comparison of Shell Matrix Proteins of four Conchifernsunder “Setting 1”. Setting 1 was set the threshold of ≥50% sequence homology, and e-value of ≤e-5

**Supplementary Table 2.**
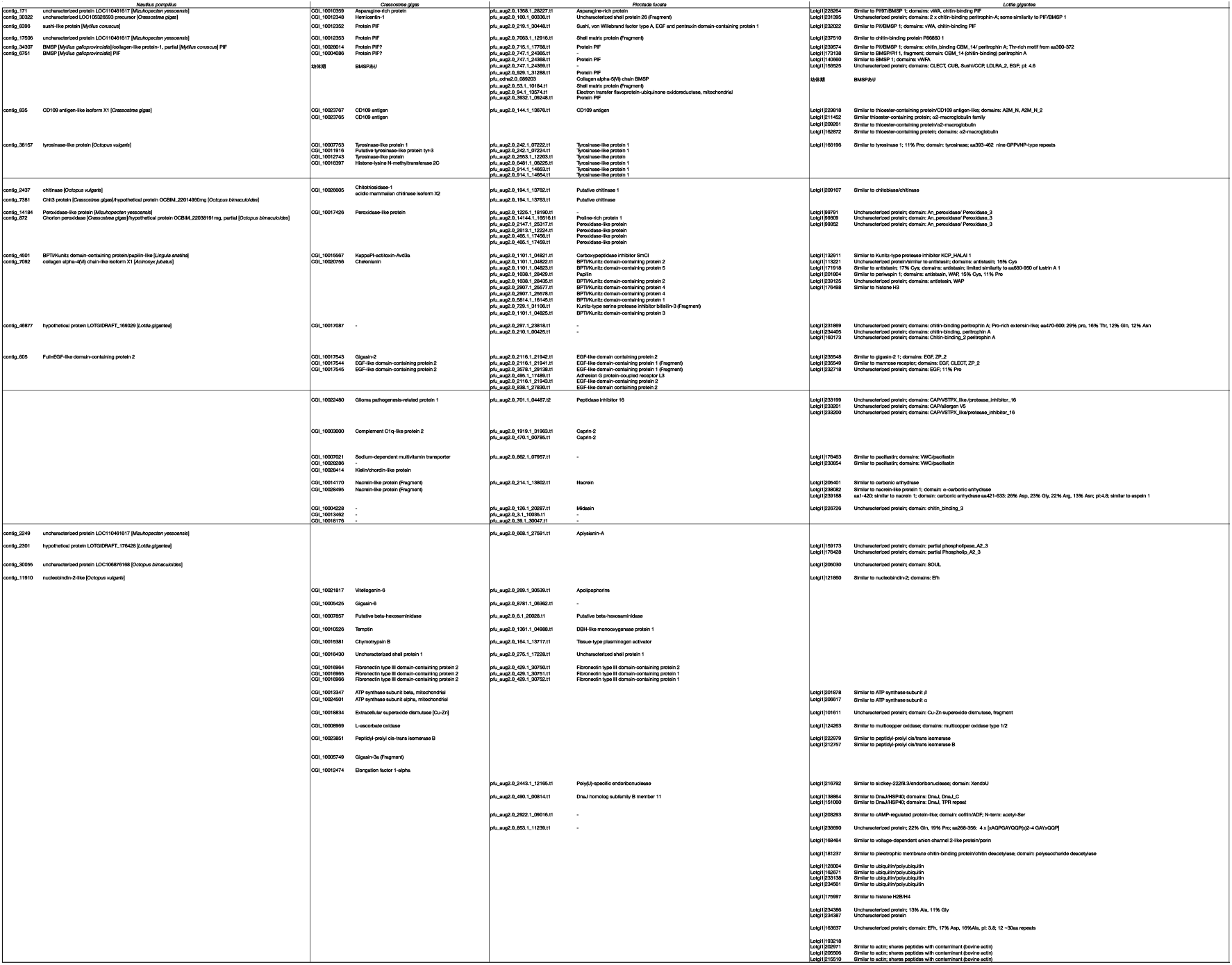
Comparison of Shell Matrix Proteins of 4 Conchiferans under “Search Setting 2”. Search Setting 2 was set the threshold of e-value of ≤e-5

**Supplementary Table 3.**
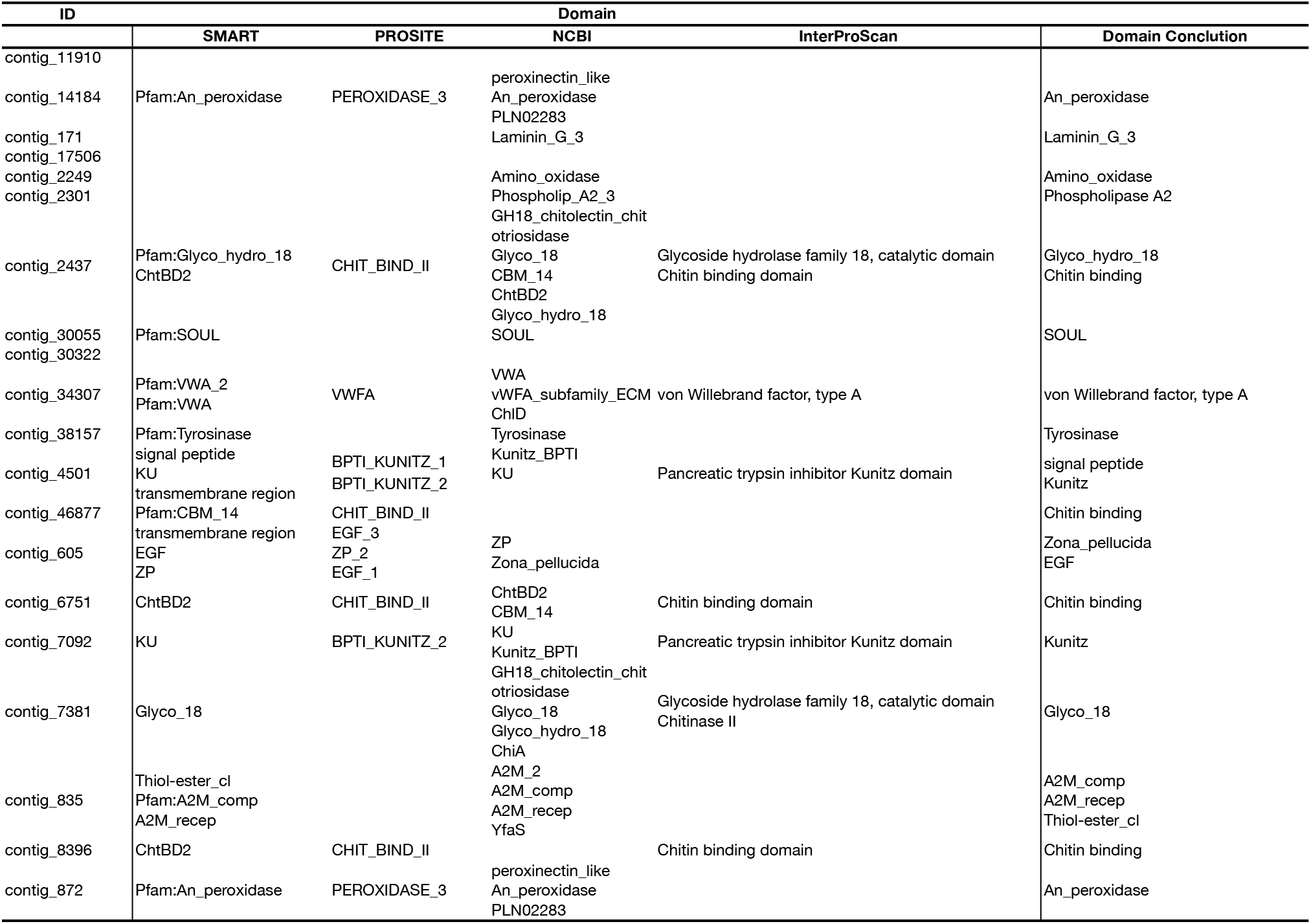
The domains of *Nautilus pompilius* as predicted by SMART, PROSITE NCBI and InterProScan

**Supplementary Table 4.**
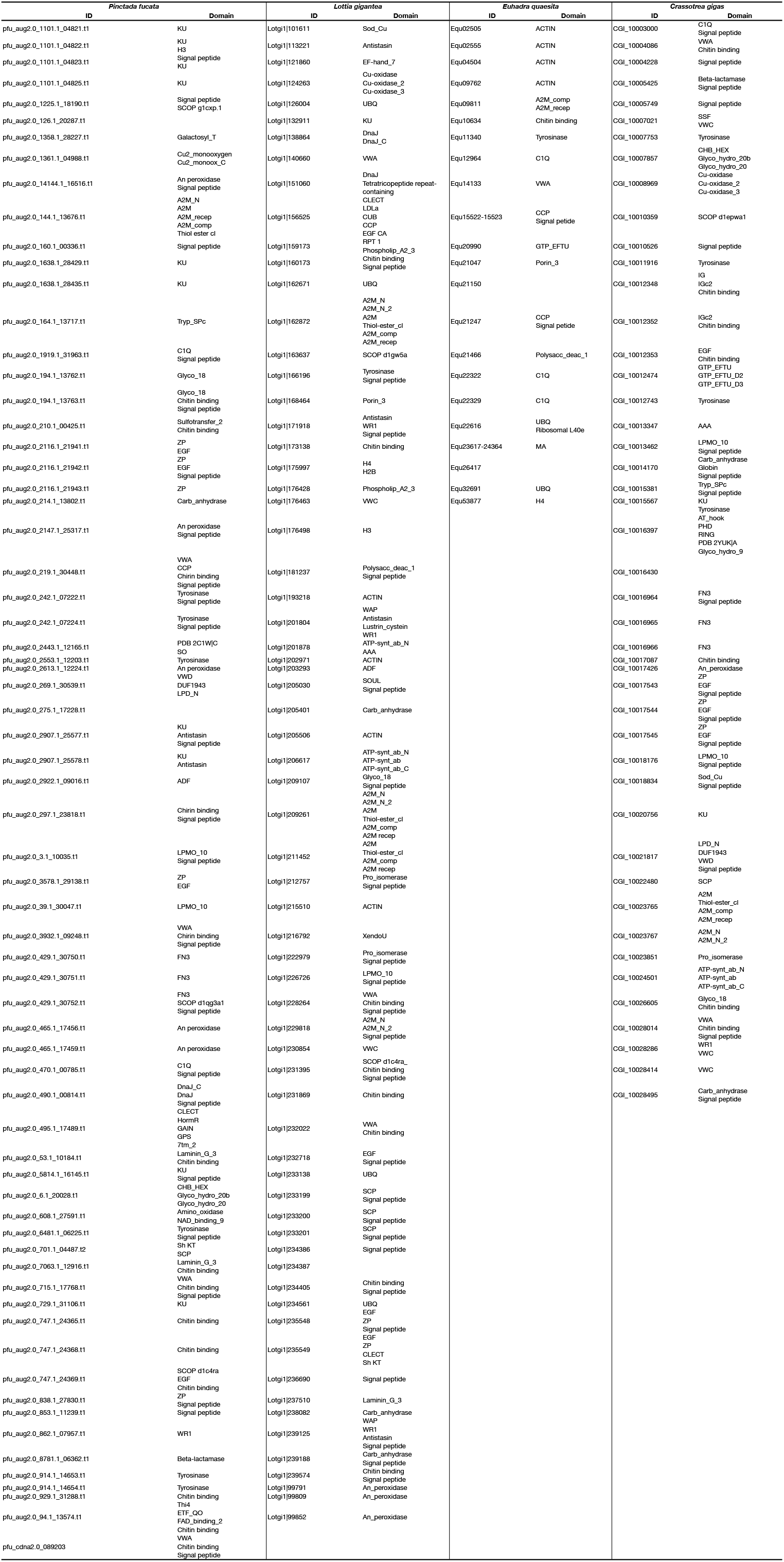
The domain of 4 spesies (*Pinctada fucata, Lottia gigantea, Euhadra quaesita*, and *Crassostrea gigas*) as predicted by SMART.

**Supplementary Table 5.** Comparison of the conserved domains of 5 species (*Nautilus pompilius, Pinctada fucata, Lottia gigantea, Euhadra quaesita*, and *Crassostrea gigas*) in Conchifera

| *N. po* | *P. fu* | *L. gi* | *E. qu* | *C. gi* |
| --- | --- | --- | --- | --- |
| A2M_comp | A2M_comp | A2M_comp | A2M_comp | A2M_comp |
| A2M_recep | A2M_recep | A2M_recep | A2M_recep | A2M_recep |
| Chitin binding | Chitin binding | Chitin binding | Chitin binding | Chitin binding |
| signal peptide | signal peptide | signal peptide | signal peptide | signal peptide |
| Tyrosinase | Tyrosinase | Tyrosinase | Tyrosinase | Tyrosinase |
| VWA | VWA | VWA | VWA | VWA |
| ZP | ZP | ZP |  | ZP |
| KU | KU | KU |  | KU |
| EGF | EGF | EGF |  | EGF |
| An_peroxidase | An_peroxidase | An_peroxidase |  | An_peroxidase |
| Glyco_18 | Glyco_18 | Glyco_18 |  | Glyco_18 |
| Thiol-ester_cl | Thiol ester cl | Thiol-ester_cl |  | Thiol-ester_cl |
| Laminin_G_3 | Laminin_G_3 | Laminin_G_3 |  |  |
| Amino_oxidase | Amino_oxidase |  |  |  |
| Phospholip_A2_3 |  | Phospholip_A2_3 |  |  |
| SOUL |  | SOUL |  |  |
|  | CCP | CCP | CCP |  |
|  | A2M | A2M |  | A2M |
|  | A2M_N | A2M_N |  | A2M_N |
|  | Carb_anhydrase | Carb_anhydrase |  | Carb_anhydrase |
|  | LPMO_10 | LPMO_10 |  | LPMO_10 |
|  | SCP | SCP |  | SCP |
|  | WR1 | WR1 |  | WR1 |
|  | C1Q |  | C1Q | C1Q |
|  | ADF | ADF |  |  |
|  | Antistasin | Antistasin |  |  |
|  | CLECT | CLECT |  |  |
|  | DnaJ | DnaJ |  |  |
|  | DnaJ_C | DnaJ_C |  |  |
|  | H3 | H3 |  |  |
|  | Sh KT | Sh KT |  |  |
|  | Beta-lactamase |  |  | Beta-lactamase |
|  | CHB_HEX |  |  | CHB_HEX |
|  | DUF1943 |  |  | DUF1943 |
|  | FN3 |  |  | FN3 |
|  | Glyco_hydro_20 |  |  | Glyco_hydro_20 |
|  | Glyco_hydro_20b |  |  | Glyco_hydro_20b |
|  | LPD_N |  |  | LPD_N |
|  | Tryp_SPc |  |  | Tryp_SPc |
|  | VWD |  |  | VWD |
|  |  | ACTIN | ACTIN |  |
|  |  | H4 | H4 |  |
|  |  | Polysacc_deac_1 | Polysacc_deac_1 |  |
|  |  | Porin_3 | Porin_3 |  |
|  |  | UBQ | UBQ |  |
|  |  | A2M_N_2 |  | A2M_N_2 |
|  |  | AAA |  | AAA |
|  |  | ATP-synt_ab |  | ATP-synt_ab |
|  |  | ATP-synt_ab_C |  | ATP-synt_ab_C |
|  |  | ATP-synt_ab_N |  | ATP-synt_ab_N |
|  |  | Cu-oxidase |  | Cu-oxidase |
|  |  | Cu-oxidase_2 |  | Cu-oxidase_2 |
|  |  | Cu-oxidase_3 |  | Cu-oxidase_3 |
|  |  | Pro_isomerase |  | Pro_isomerase |
|  |  | Sod_Cu |  | Sod_Cu |
|  |  | VWC |  | VWC |
|  |  |  | GTP_EFTU | GTP_EFTU |

**Supplementary Table 6.** The specific domains of 5 species (*Nautilus pompilius, Pinctada fucata, Lottia gigantea, Euhadra quaesita*, and *Crassostrea gigas*) in Conchifera

| Npo | Pfu | Lgi | Equ | Cgi |
| --- | --- | --- | --- | --- |
| Glyco_hydro_18 | Cu2_monoox_C | CUB | MA | AT_hook |
|  | 7tm_2 | EF-hand_7 | Ribosomal<br>L40e | Globin |
|  | Cu2_monooxygen | EGF CA |  | Glyco_hydro_9 |
|  | ETF_QO | H2B |  | GTP_EFTU_D2 |
|  | FAD_binding_2 | LDLa |  | GTP_EFTU_D3 |
|  | GAIN | Lustrin_cystein |  | IG |
|  | Galactosyl_T | RPT 1 |  | IGc2 |
|  | GPS | SCOP d1c4ra_ |  | PDB 2YUK\|A |
|  | HormR | SCOP d1gw5a |  | PHD |
|  | NAD_binding_9 | Tetratricopeptide repeat-containing domain |  | RING |
|  | PDB 2C1W\|C |  |  | SCOP d1epwa1 |
|  | SCOP d1c4ra |  |  | SSF |
|  | SCOP d1qg3a1 |  |  |  |
|  | SCOP g1cxp.1 |  |  |  |
|  | SO |  |  |  |
|  | Sulfotransfer_2 |  |  |  |
|  | Thi4 |  |  |  |

